## Supplementary Data for "New structural features of the APC/C from high-resolution cryo-EM structures of apo-APC/C and APC/C^CDH1:EMI1^ complexes"

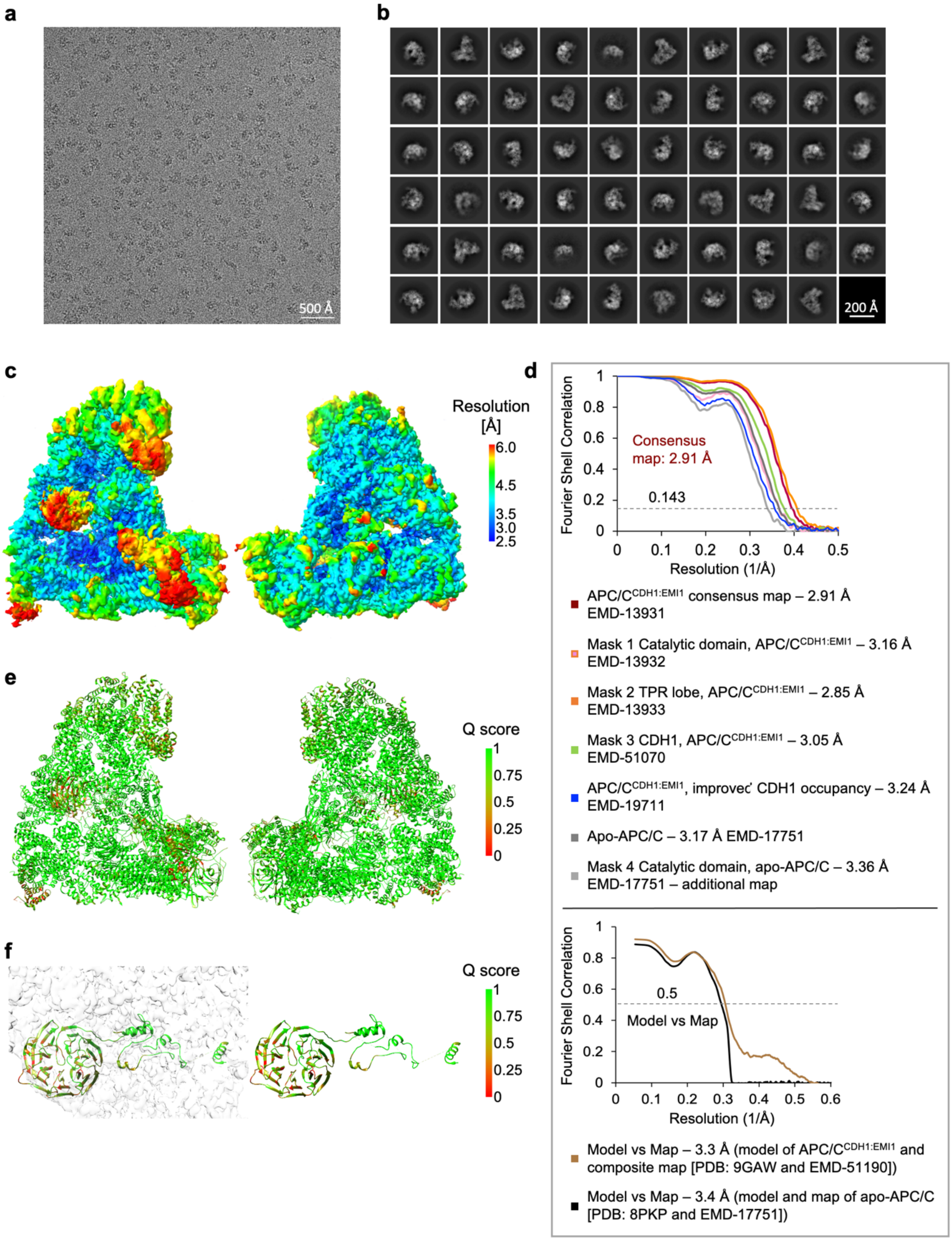


**Supplementary Figure 1. Cryo-EM images and refined APC/C reconstructions. a**, Representative cryo-EM image of the $APC/C^{CDH1:EMI1}$ complex. **b**, Gallery of 2D-class averages of $APC/C^{CDH1:EMI1}$. **c**, Local resolution map of $APC/C^{CDH1:EMI1}$. **d**, Fourier Shell Correlation (FSC) plots of the main maps used to generate molecular models. **e**, Q-score analysis [43] of the $APC/C^{CDH1:EMI1}$ complex to validate map-model correlation. **f**, Close-up view on the CDH1 subunit of the $APC/C^{CDH1:EMI1}$ complex that is colour-coded by Q-score.

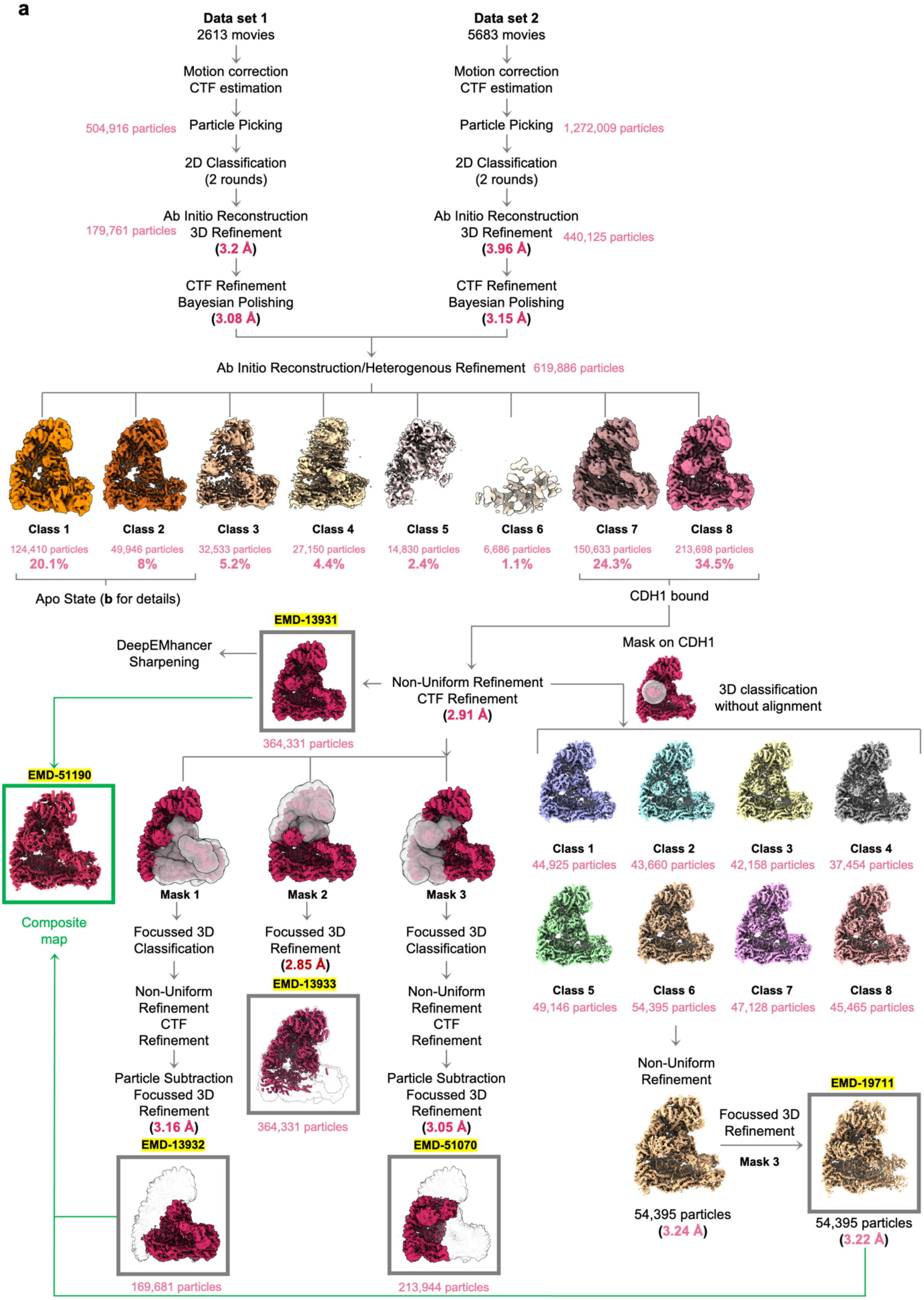
a
Data set 1
2613 movies
Motion correction
CTF estimation
504,916 particles
Particle Picking
2D Classification
(2 rounds)
Ab Initio Reconstruction
179,761 particles
3D Refinement
(3.2 Å)
CTF Refinement
Bayesian Polishing
(3.08 Å)
Data set 2
5683 movies
Motion correction
CTF estimation
Particle Picking
1,272,009 particles
2D Classification
(2 rounds)
Ab Initio Reconstruction
3D Refinement
440,125 particles
(3.96 Å)
CTF Refinement
Bayesian Polishing
(3.15 Å)
Ab Initio Reconstruction/Heterogenous Refinement 619,886 particles
Class 1
124,410 particles
20.1%
Class 2
49,946 particles
8%
Class 3
32,533 particles
5.2%
Class 4
27,150 particles
4.4%
Class 5
14,830 particles
2.4%
Class 6
6,686 particles
1.1%
Class 7
150,633 particles
24.3%
Class 8
213,698 particles
34.5%
Apo State (b for details)
CDH1 bound
EMD-13931
DeepEMhancer
Sharpening
Mask on CDH1
Non-Uniform Refinement
CTF Refinement
(2.91 Å)
3D classification
without alignment
364,331 particles
EMD-51190
Composite
map
Mask 1
Focussed 3D
Classification
Non-Uniform
Refinement
CTF
Refinement
Particle Subtraction
Focussed 3D
Refinement
(3.16 Å)
EMD-13932
169,681 particles
Mask 2
Focussed 3D
Refinement
(2.85 Å)
EMD-13933
364,331 particles
Mask 3
Focussed 3D
Classification
Non-Uniform
Refinement
CTF
Refinement
Particle Subtraction
Focussed 3D
Refinement
(3.05 Å)
EMD-51070
213,944 particles
Class 1
44,925 particles
Class 2
43,660 particles
Class 3
42,158 particles
Class 4
37,454 particles
Class 5
49,146 particles
Class 6
54,395 particles
Class 7
47,128 particles
Class 8
45,465 particles
Non-Uniform
Refinement
54,395 particles
(3.24 Å)
Focussed 3D
Refinement
Mask 3
EMD-19711
54,395 particles
(3.22 Å)

**b**

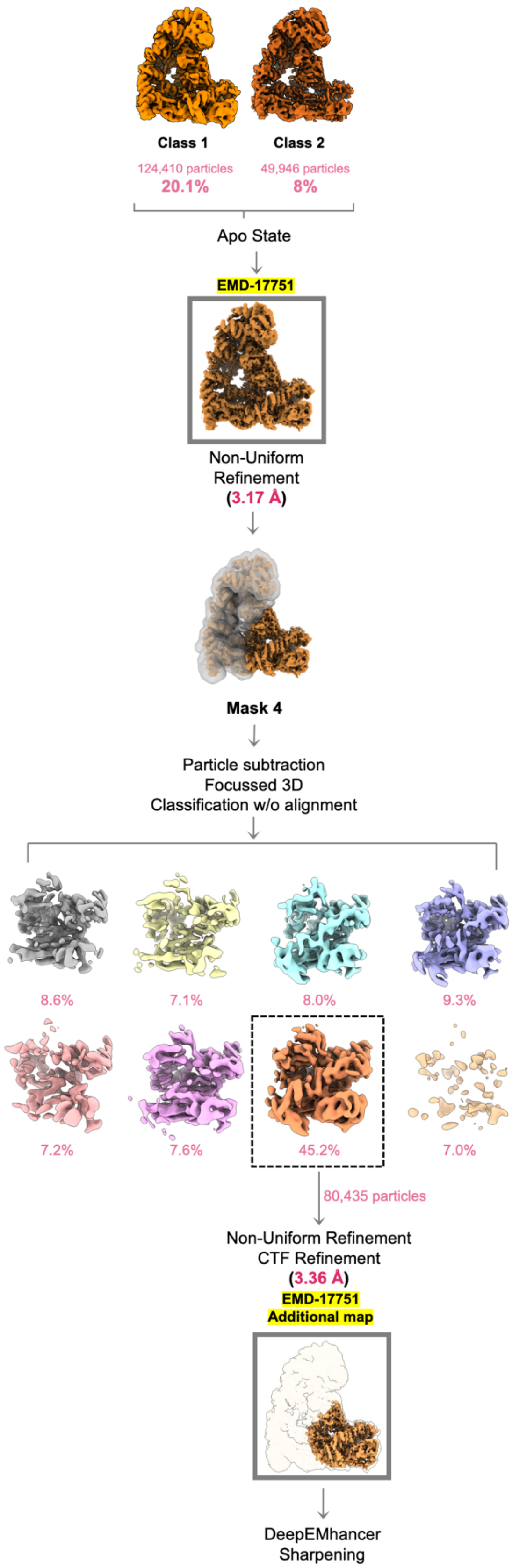


**Supplementary Figure 2. Data processing pipeline for cryo-EM reconstructions. (a)** Cryo-EM data processing workflow summary as described in the Methods section for the ternary APC/C$^{CDH1:EMI}$ complex. (**b**) Work flow for apo-APC/C.

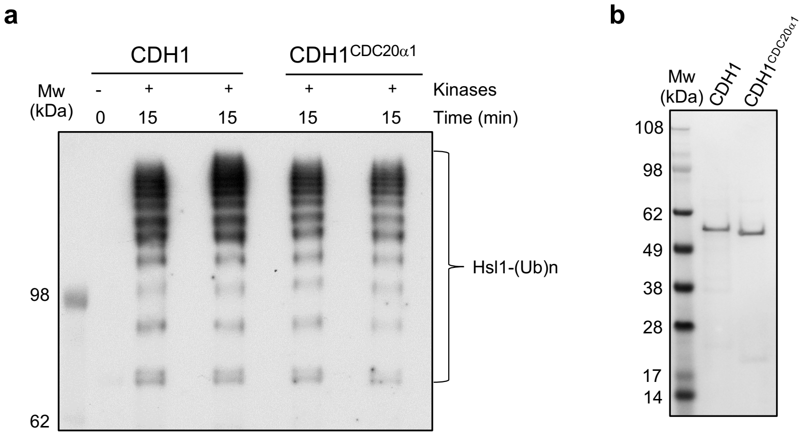


**Supplementary Figure 3. CDH1 and CDH1$^{CDC20\alpha1}$ activate unphosphorylated and phosphorylated APC/C to similar extents. a**, Ubiquitination assay with unphosphorylated and phosphorylated APC/C activated by CDH1 and CDH1$^{CDC20\alpha1}$. **b**, SDS PAGE gel of CDH1 and CDH1$^{CDC20\alpha1}$ used in the ubiquitination assay.

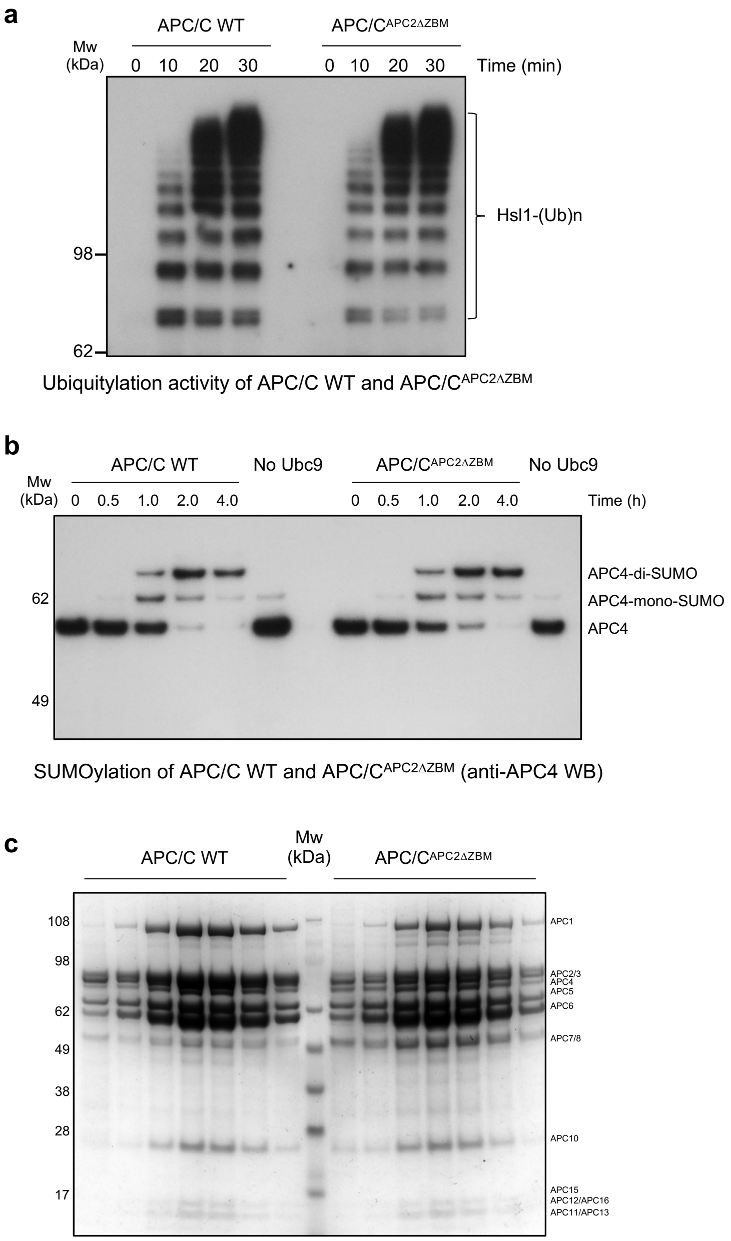


**Supplementary Figure 4. Wild type APC/C and the APC/C$^{APC2\Delta ZBM}$ mutant have similar ubiquitination activity and are SUMOylated to the same extent. a**, A ubiquitylation assay shows wild type APC/C and the APC/C$^{APC2\Delta ZBM}$ mutant (disrupted APC2$^{ZBM}$) are equally active as E3 ligases. **b**, SUMOylation assays shows that wild type APC/C and the APC/C$^{APC2\Delta ZBM}$ mutant are SUMOylated to the same extent. APC4 SUMOylation is shown by a Western blot against APC4. **c**, SDS PAGE gel comparing wild type APC/C and mutant APC/C$^{APC2\Delta ZBM}$. The mutant APC2$^{\Delta ZBM}$ did not impair APC/C assembly.

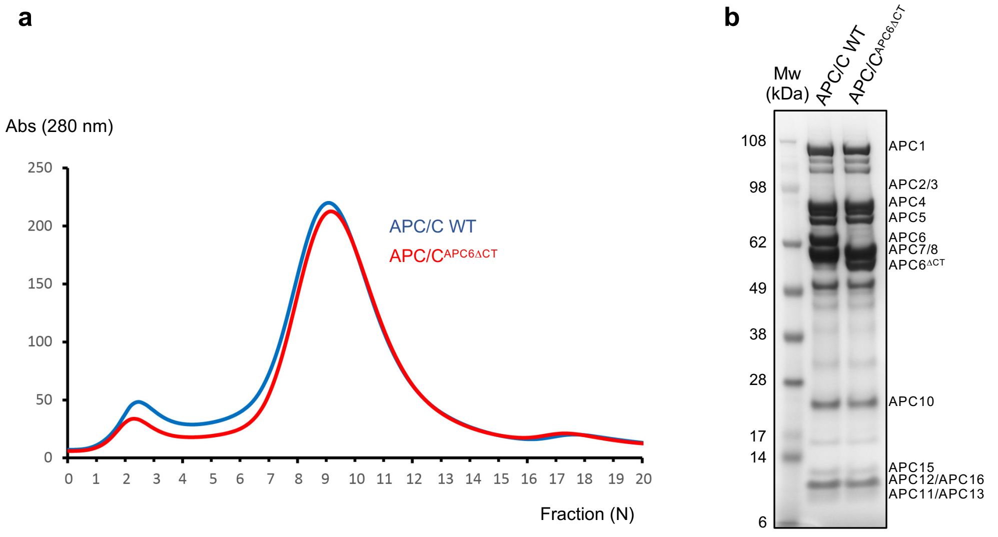


SDS PAGE gel of wild type APC/C and APC/C$^{\Delta APC6CT}$

**Supplementary Figure 5. APC/C$^{APC6\Delta CT}$ assembles normally. a,** Size exclusion chromatogram and **b**, SDS PAGE gel of wild type APC/C and APC/C$^{APC6\Delta CT}$ show similar elution profiles on a SEC column (a), and subunit stoichiometry (b).

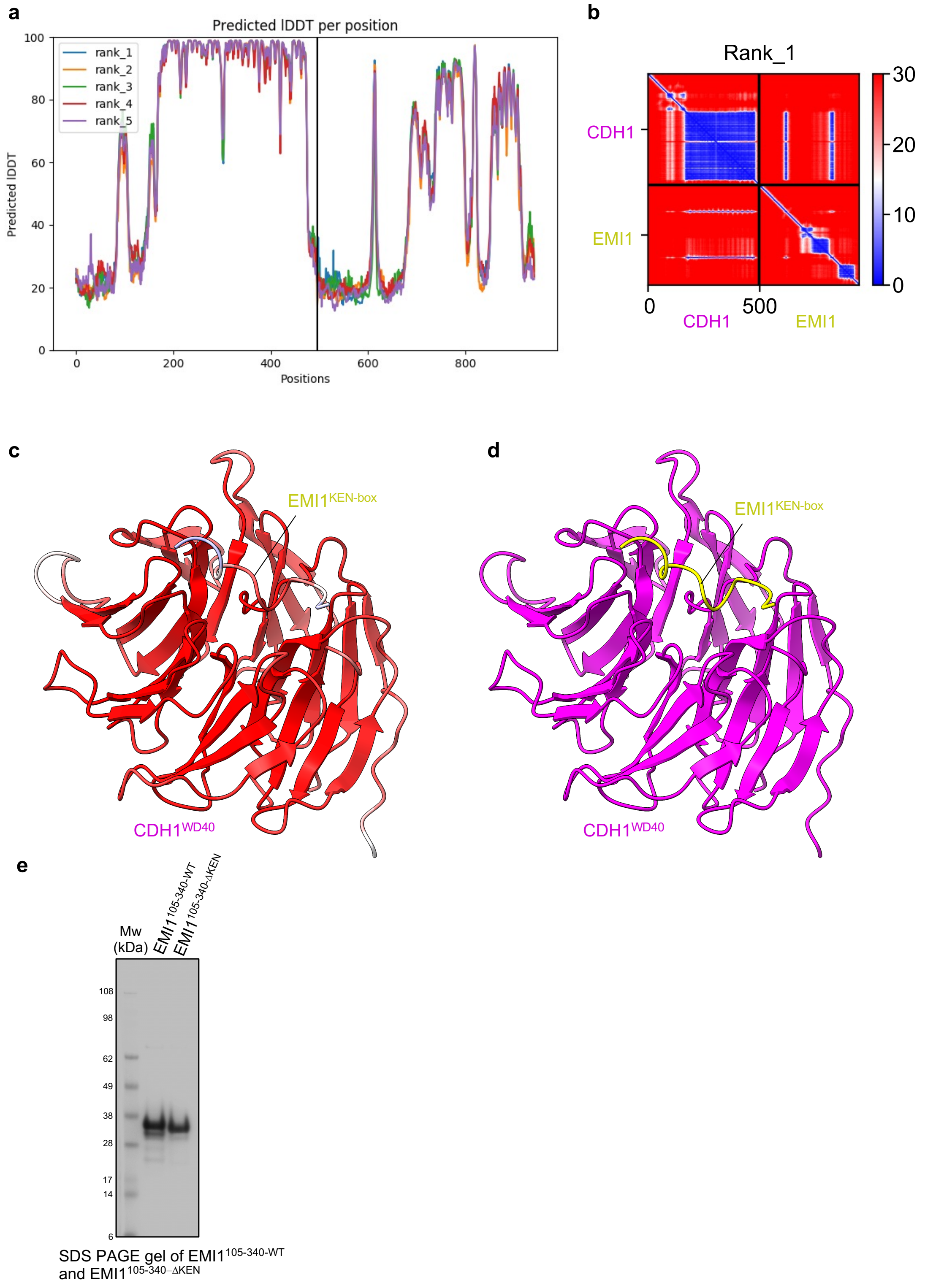


**Supplementary Figure 6. AlphaFold2 prediction of the CDH1$^{WD40}$:EMI1$^{KEN\ box}$ interaction and confidence metrics**. **a**, Predicted local distance difference test (lDDT) values for the AlphaFold2 models of CDH1$^{WD40}$:EMI1$^{KEN\ box}$. A higher score indicates higher confidence in the prediction. **b**, The predicted alignment error (PAE) heat map for the CDH1$^{WD40}$:EMI1$^{KEN\ box}$ AlphaFold2 prediction. The PAE heat map shows the predicted error (in angstroms) between all pairs of residues, with blue indicating lower error and red indicating higher error. The PAE plot suggests high confidence for the predicted interactions between CDH1$^{WD40}$ and EMI1$^{KEN\ box}$. **c**, The pLDDT values mapped onto the predicted model of CDH1$^{WD40}$:EMI1$^{KEN\ box}$. Residues are coloured in a scale of high (red) to low (blue) pLDDT values. The model was predicted with high confidence (pLDDT > 90). **d**, Predicted model colour-coded according to Fig. 1c. **e**, SDS PAGE gel of EMI1$^{105\text{-}340\text{-}WT}$ and EMI1$^{105\text{-}340\text{-}\Delta KEN}$.

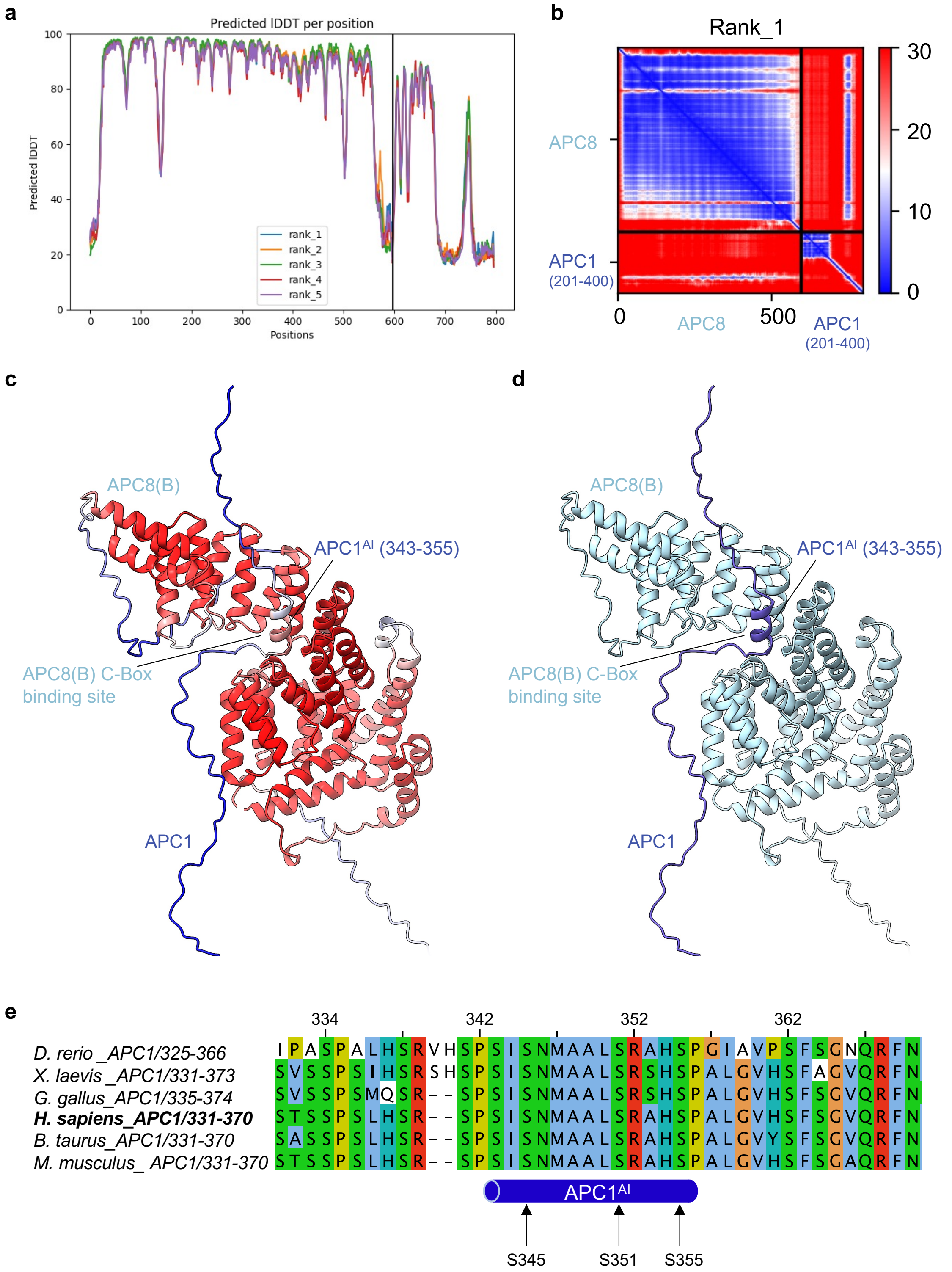


**Supplementary Figure 7. AlphaFold2 prediction of the APC1$^{AI}$:APC8 interaction and confidence metrics**. **a**, Predicted local distance difference test (lDDT) values for the AlphaFold2 models of APC1$^{AI}$:APC8. A higher score indicates higher confidence in the prediction. **b**, The predicted alignment error (PAE) heat map for the APC1$^{AI}$:APC8 AlphaFold2 prediction. The PAE heat map shows the predicted error (in angstroms) between all pairs of residues, with blue indicating lower error and red indicating higher error. The PAE plot suggests high confidence for the predicted interactions between APC1$^{AI}$ and APC8. **c**, The pLDDT values mapped onto the predicted model of APC1$^{AI}$:APC8. Residues are coloured in a scale of high (red) to low (blue) pLDDT values. The model was predicted with high confidence (pLDDT > 90). **d**, Predicted model colour-coded according to Fig. 1c. **e**, MSA of APC1 comprising the AI segment of selected vertebrate species. Sites of phosphoryation are indicated (Ser345, Ser351, Ser355).

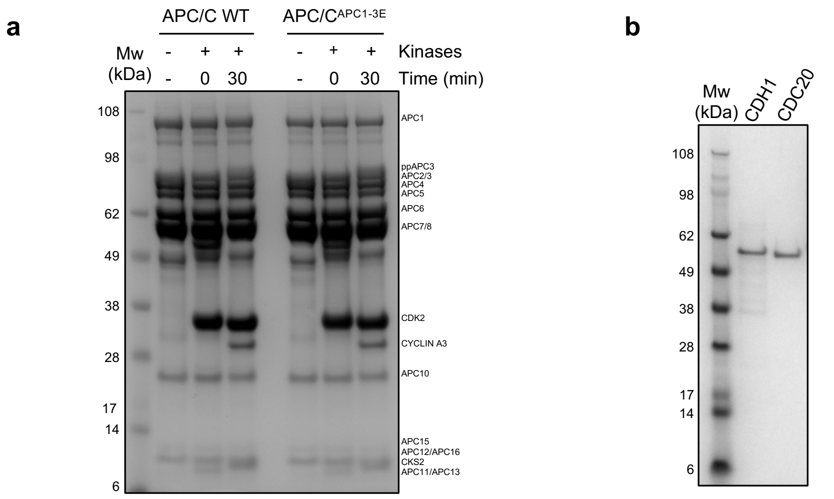


SDS PAGE gel of wild type APC/C and APC/C$^{APC1\text{-}3E}$ before and after phosphorylation

SDS PAGE gel of CDH1 and CDC20

**Supplementary Figure 8. SDS PAGE gels of purified APC/C complexes and coactivator**. **a**, SDS PAGE gel of unphosphorylated and phosphorylated wild type APC/C and APC/C$^{APC1\text{-}3E}$ mutant. The slower migrating phosphorylated APC3 subunit (ppAPC3) indicates APC/C phosphorylation. **b**, SDS PAGE gel of CDH1 and CDC20 used in this assay.

### Supplementary Table 1. Cryo-EM data collection, refinement and validation statistics

| | | APC/C$^{CDH1:EMI1}$ Dataset 1 | | | APC/C$^{CDH1:EMI1}$ Dataset 2 | | |
|---|---|---|---|---|---|---|---|
| **Data collection** | | | | | | | |
| Microscope | | FEI Titan Krios | | | FEI Titan Krios | | |
| Voltage (keV) | | 300 | | | 300 | | |
| Electron dose ($e^-$/Å$^{-2}$) | | 40 | | | 40 | | |
| Detector | | FEI Falcon III | | | FEI Falcon III | | |
| Pixel size (Å/pixel) | | 1.07 | | | 1.07 | | |
| Defocus range (μm) | | 1.5-3.5 | | | 1.5-3.5 | | |
| Micrographs (N) | | 2,613 | | | 5,683 | | |
| **Reconstructions** | APC/C$^{apo}$ | APC/C$^{CDH1:EMI1}$ Composite map | APC/C$^{CDH1:EMI1}$ Consensus map | APC/C$^{CDH1:EMI1}$ Focussed Map Catalytic domain<br><br>Mask1 | APC/C$^{CDH1:EMI1}$ Focussed Map TPR lobe<br><br>Mask2 | APC/C$^{CDH1:EMI1}$ Focussed Map CDH1<br><br>Mask3 | APC/C$^{CDH1:EMI1}$ Improved CDH1 occupancy |
| Particles (N) | 174,356 | - | 364,331 | 169,681 | 364,331 | 213,944 | 54,395 |
| Box size (pix) | 360 | 360 | 360 | 360 | 360 | 360 | 360 |
| Map sharpening | | | | - | | - | |
| B-factor (Å$^2$) | -116.8 | - | -119.1 | 106.3 | -117.3 | 121.3 | -118.5 |
| Resolution (global, Å) | 3.2 | - | 2.9 | 3.16 | 2.85 | 3.05 | 3.24 |
| Resolution range (local, Å) | 2.7-9.1 | - | 2.4-7.8 | 2.6-9.0 | 2.4-8.5 | 2.6-8.6 | 2.7-9.5 |
| FSC threshold | 0.143 | - | 0.143 | 0.143 | 0.143 | 0.143 | 0.143 |
| **Model composition** | | | | | | | |
| Protein residues (N) | 8,076 | 8,600 | | | | | |
| Zn ions (N) | 4 | 6 | | | | | |
| **Refinement** | | | | | | | |
| Resolution (Å) | 3.4 | 3.3 | | | | | |
| FSC threshold | 0.5 | 0.5 | | | | | |
| Model to map (CC) | 0.78 | 0.81 | | | | | |
| *B* factors (Å$^2$) | | | | | | | |
| Protein residues | -79.1 | -62.9 | | | | | |
| Ligands | -205.5 | -184.5 | | | | | |
| RMS deviation | | | | | | | |
| Bond lengths (Å) | 0.004 | 0.005 | | | | | |
| Bong angles (°) | 0.642 | 1.008 | | | | | |
| **Validation** | | | | | | | |
| Clashscore, all atoms | 12 | 11 | | | | | |
| Rotamer outliers (%) | 2.3 | 0.8 | | | | | |
| **Ramachandran plot** | | | | | | | |
| Favoured (%) | 97.4 | 96.9 | | | | | |
| Allowed (%) | 2.60 | 3.11 | | | | | |
| Outliers (%) | 0 | 0.02 | | | | | |
| **Deposition** | | | | | | | |
| PDB ID | 8PKP | 9GAW | | | | | |
| EMDB ID | EMD-17751 | EMD-51190 | EMD-13931 | EMD-13932 | EMD-13933 | EMD-51070 | EMD-19711 |

**Supplementary Table 2. Rebuilt regions of APC/C$^{CDH1:EMI1}$ subunits PDB: 9GAW**

| Subunit | Residue range | Modification | Comments | 4ul9 |
|---|---|---|---|---|
| **APC1** | 135-145 | Loop refitted | | |
| | 192-206 | Loop refitted | | |
| | 275-283 | Rebuild | | |
| | 282-295 | α-helix to loop<br>Helix now Cdh1$^N$ | | |
| | 415-424 | Loop fitted | | |
| | 460-464 | Loop fitted | | |
| | 646-650 | Loop rebuilt | | |
| | 671-681 | α-helix built | | |
| | 701-734 | Build | | SC unassigned |
| | 741-757 | Build | | |
| | 814-838 | Build | | |
| | 853-859 | Build | | |
| | 882-901 | Build | | |
| | 1009-1013 | Build | | |
| | 1333-1347 | Build | | |
| | 1433-1452 | Build | | |
| | 1673-1684 | Rebuild | | |
| | 1711-1734 | Rebuild/build | | |
| | 1734-1911 | Renumber -3 | | |
| | 1825-1839 | Build | | |
| | 1873-1877 | Build | | |
| | 1897-1911 | Build | | |
| | 1897-1936 | Rebuild | | |
| | 1911-1935 | Reverse chain polarity, fit s.c. | | |
| **APC2** | 17-35 | s.c. built to α-helix | | 15-28, m.c. only |
| | 46-52 | Build | | |
| | 53-87 | Rebuild and fit s.c. | | |
| | 126-147 | Fit s.c. | | Poly A |
| | 187-233 | Build/rebuild, fit s.c. | New Zn-binding module built: Cys 221,224, 231,233 | |
| | 304-322 | Loop fitted | | |
| | 458-507 | Rebuild | | |
| | 605-640 (CTD) | | Good density | |
| | 644-659 (CTD) | | β-hairpin no density | |
| | 660-716 (CTD) | m.c. good, s.c. poorly defined | Only poor part of APC2 | |
| **APC3(B) chain P** | 171-176 | Build | | |
| | 177-445 | | Long disordered region | |
| | 446-450 | Build | | |
| **APC3(A) chain J** | 2-4 | Build | | |
| | 171-176 | Build | | |
| | 177-445 | | Long disordered region | |
| | 446-450 | Build | | |
| | 767-780 | Rebuild | | |
| **APC4** | 126-133 | Loop fitted | | |
| | 276-301 | Rebuild | | |
| | 430-439 | Loop fitted | | |

| | | | | |
|---|---|---|---|---|
| | 465-470 | Loop fitted | | |
| | 487-495 | Rebuild | | |
| **APC5** | 9-27 | Build | | |
| | 46-55 | Rebuild | | |
| | 351-357 | Rebuild | | |
| | 742-755 | Rebuild | | |
| **APC6(A) chain Q** | No change | | | |
| **APC6(B) chain K** | 94-98 | Build | | |
| | 120-128 | Build | | |
| | 544-567 | Build: fit s.c. assigned to APC6 | | Poly A, unassigned chain |
| **APC7(A) chain Y** | 34-35 | Build | | |
| | 553-555 | Build | | |
| **APC7(B) chain Z** | 34-35 | Build | | |
| | 131 | Build | | |
| | 111-122 | Build | | |
| | 371-540 | Reposition $\alpha$-helices | | |
| **APC8(A) chain U** | 499-509 | Loop fitted | | |
| | 504-523 | Rebuild | | |
| **APC8(B) chain V** | 499-509 | Loop fitted | | |
| **APC10** | 167-176 | Rebuild, s.c. fitted | | |
| **APC11** | 1-8 | Build $\beta$-strand | | |
| | 18-84 | RING domain | m.c. and s.c.: good fit | |
| **APC12(A) chain G** | 26-27 | Build | | |
| **APC12(B) chain W** | 26 | Build | | |
| **APC13** | 1-6 | Rebuild | | |
| | 39-47 | Build | | |
| | 68 | Build | | |
| **APC15** | No change | | | |
| **APC16** | 51-57 | Rebuild | | |
| | 108 | Build | | |
| **CDH1** | 1-17 | Build as $\alpha$-helix | | $\alpha$-helix assigned to APC1 284-295 |
| | 144-145 | Build | | |
| | 164-172 | Build | Connects to WD40 | |
| | 473-476 | Rebuild | | |
| **EMI1** | 117-121 | Build | KEN-box | |
| | | ZBR domain 356-415 | m.c. and s.c.: good fit | |
| | 433-447 | Fit s.c. | Incudes LRRL | |

**Supplementary Table 3. Ordered and disordered regions of APC/C subunits**

| Subunit | Visible N-term | Visible C-term | Disordered regions | [6]AF2 C-term helix | Protein Length (N) |
|---|---|---|---|---|---|
| **APC1** | 10 | 1936 | 51-70, 195-204, 228-233, 284-399 (AI segment), 516-581, 682-700, 735-741, 902-921, 987-1010, 1336-1346, 1439-1451, 1902-1907, 1937-1944 | | 1944 |
| **APC2** | 17 | 716 | 35-48, 307-317, 461-474, 463-470, 488-495, 646-657, 707-711, 717-822 (WHB domain) | | 822 |
| **[1,2]APC3(A) chain J** | 3 | 780 | 172-450, 781-824 | 783 | 824 |
| **[1,2]APC3(B) chain P** | 5 | 769 | 174-447, 770-824 | 783 | 824 |
| **APC4** | 3 | 755 | 128-133, 431-437, 458-468, 756-808 | | 808 |
| **APC5** | 9 | 755 | 20-25, 168-204 (linker), 453-457 | | 755 |
| **[3,4]APC6(A) chain Q** | 1 | 533 | 96-123 (AF2 $\alpha$-helix 99-109), 534-630 | 528 | 620 |
| **[3]APC6(B) chain K** | 1 | 565 | 99-126 (AF2 $\alpha$-helix 99-109), 528-544, 566-620 | 528 | 620 |
| **APC7(A) chain Y** | 35 | 540 | 111-131, 541-565, AF2 $\alpha$-helix 3-23 not visible | 557 | 565 |
| **APC7(B) chain Z** | 35 | 507 | 111-130, 508-565, AF2 $\alpha$-helix 3-23 not visible | 557 | 565 |
| **[5]APC8(A) chain U** | 25 | 523 | 136-144, 524-59 | 559 | 597 |
| **[5]APC8(B) chain V** | 24 | 523 | 523-597 | 559 | 597 |
| **APC10** | 4 | 185 | 164-167 | | 185 |
| **APC11** | 3 | 83 | 84 | | 84 |
| **APC12(A) chain W** | 1 | 26 | 27-85 | | 85 |
| **APC12(B) chain G** | 1 | 26 | 27-85 | | 85 |
| **APC13** | 1 | 68 | 42-45, 69-74 | | 74 |
| **APC15** | 2 | 57 | 58-121 | | 121 |
| **APC16** | 51 | 108 | 1-50, 109-110 | | 110 |
| **CDH1** | 1 | 496 | 14-41, 68-87, 110-122,134-142, 416-434 | | 496 |
| **EMI1** | 116 | 447 | 1-115, 123-319, 335-357, 416-434 | | 447 |

Notes

[1]Different trajectory N-term of IR tail for CDH1 and APC10
[2]Residues 273-286 of APC3 insert predicted to bind IR-tail binding site of APC3, includes $T^{279}P$
[3]AF2: In free APC6, residues 596-620 of APC6 mimic residues 1-27 of APC12
[4]Residues 529-540 of APC6(A) chain Q interacts with APC8(A) chain U
[5]AF2: Residues 587-C-term of APC8 binds to C-box/IR tail site of APC8
[6]C-terminal residue of an AF2-predicted $\alpha$-helix.

N-term and C-term residues of sequence
